# *Diviner* uncovers hundreds of novel human (and other) exons though comparative analysis of proteins

**DOI:** 10.1101/2024.05.05.592595

**Authors:** Alexander J Nord, Travis J Wheeler

## Abstract

**Background:** Eukaryotic genes are often composed of multiple exons that are stitched together by *splicing* out the intervening introns. These exons may be conditionally joined in different combinations to produce a collection of related, but distinct, mRNA transcripts. For protein-coding genes, these products of *alternative splicing* lead to production of related protein variants (*isoforms*) of a gene. Complete labeling of the protein-coding content of a eukaryotic genome requires discovery of mRNA encoding all isoforms, but it is impractical to enumerate all possible combinations of tissue, developmental stage, and environmental context; as a result, many true exons go unlabeled in genome annotations.

**Results:** One way to address the combinatoric challenge of finding all isoforms in a single organism *A* is to leverage sequencing efforts for other organisms – each time a new organism is sequenced, it may be under a new combination of conditions, so that a previously unobserved isoform may be sequenced. We present *Diviner*, a software tool that identifies previously undocumented exons in organisms by comparing isoforms across species. We demonstrate *Diviner*’s utility by locating hundreds of novel exons in the genomes of human, mouse, and rat, as well as in the ferret genome. Further, we provide analyses supporting the notion that most of the new exons reported by *Diviner* are likely to be part of a true (but unobserved) isoform of the containing species.

## Introduction

When identifying the protein-coding regions of eukaryotic genomes, a challenge arises from the fact that most genes are made up of multiple exons that are fused together as the non-coding introns are spliced from a pre-mRNA molecule during mRNA maturation (1). One important consequence of this mechanism is that exons can be included or excluded during mRNA production in a condition-dependant manner (Figure 1).

**Fig. 1.**
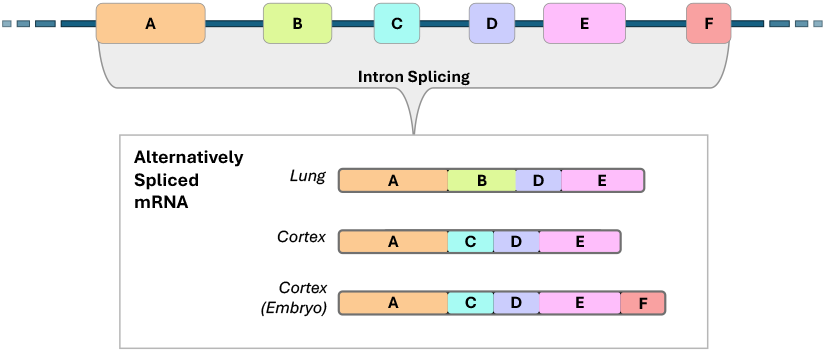
A toy diagram illustrating how alternative splicing, influenced by tissue type and environmental factors, can result in mRNA sequences constituted by distinct sets of exons. These varying mRNA encode related but different protein products, called isoforms.

Alternative splicing produces a set of related, but distinct, protein variants for a single gene, called *isoforms*; these protein variants typically have varying folding and binding behavior (2, 3), and their production is influenced by cell type, developmental stage, and environmental conditions/stimuli (4–6).

Purely computational identification of genes and their isoforms remains an open challenge (7). The most reliable method for identifying isoforms (and their constituent exons) is through experimental evidence of exon utilization (8), typified by RNA-Seq or ribosomal profiling (9, 10). Ideally, one might enumerate all combinations of conditions influencing alternative splicing, then sequence the mRNA present in each enumerated sample, and thus acquire complete insight into the diversity of isoforms and the identity of exons that make them up. In practice, however, enumeration is infeasible due to the extreme combinatorics of cell type, developmental stage, and (especially) environmental condition. Importantly, it is *a priori* unknowable whether any set of sequencing circumstances would be sufficient to induce the full spectrum of mRNA transcripts required to observe every exon that can be incorporated into a given gene. One remedy for this combinatoric obstacle is to leverage sequencing efforts across a diverse pool of organisms: each sequencing effort may explore a new set of conditions (cell type, developmental stage, environmental state) and thus identify a previously-unobserved isoform. Sequence similarity with matching splice boundaries to an observed isoform suggests evolutionary conservation of coding potential (11). We can advance annotation of the exonic landscape of genomes by identifying exons in one species that are not observed in another species, then locating the homologous unannotated exons in that other genome (12, 13).

A naïve implementation of this homology-based exon identification approach is to align each isoform observed in one species against the genomes of many target organisms using a spliced alignment tool such as *spaln2* (14) or *miniprot* (15). While this approach will enable exon annotation, it will incur significant and unnecessary computational expense (performing full-genome mapping, and identifying previously-known exons) and is subject to risk of false labeling (due to spurious similarities to windows of genomic sequence outside of a gene’s neighborhood). To improve on this stratgy: (1) annotation search should focus only on observed peptides encoded by exons in one genome without known homologs on the other genome, and (2) the search region on the target genome should be restricted to a narrow window where the presence of an exon homolog would be biologically consistent with the locations of neighboring exons.

Here, we present *Diviner*, a software tool that uses targeted sequence alignment to identify unannotated coding regions. *Diviner* uses splice-aware isoform multiple sequence alignments (MSAs) produced by our *Mirage2* (16) software to quickly identify exons that appear to lack a known homolog in one or more organisms. Using *Mirage2*’s protein-to-genome mapping coordinates, *Diviner* then performs a narrow search of the target organism’s genome precisely where a missing exon homolog is expected to be encoded. Alignment of the peptide for the apparently-missing exon to the appropriate genomic window identifies coding regions that have not yet been annotated as exons.

Tests performed on the human, mouse, and rat genomes demonstrate that *Diviner* locates 663 exon homologs that have not been previously annotated or predicted as coding regions in these model organisms’ genomes (*i*.*e*., that are “entirely novel” predictions). In addition to its use in model organisms, *Diviner*’s potential applications for annotating the genomes of less-studied organisms are illustrated by its discovery of 699 entirely novel coding regions in the ferret genome. Finally, several analyses are provided that indicate that the quantitative features of the coding regions predicted by *Diviner* agree with those of previously-observed exons, bolstering confidence in the validity of the predicted exons.

## Results

The primary result of Diviner is identification of a large collection of previously unannotated exons in human, mouse, rat, and ferret genomes. The majority of these newly identified exons are supported by high quality annotation in one species combined with reliable inter-species sequence alignment to another. We begin by briefly describing the procedure for exon discovery (see Methods for more detail), then present information about the set of discovered exons.

### Method Sketch

*Diviner*’s exon discovery process depends on inter-species multiple sequence alignments (MSAs) of protein isoforms produced by our software, *Mirage2* (16). *Mirage2* works by first mapping each isoform sequence to its coding nucleotides on the genome and then using those nucleotide mapping coordinates as the basis for intra-species alignment of isoforms; subsequent alignment of isoforms across species depends on a scoring scheme that rewards splice site concordance. *Diviner* ingests *Mirage2* MSAs and collapses the sequences of all isoforms within a species into a single sequence representing the full ordered set of exons in the gene, resulting in an MSA with one *all-exon* sequence per organism. When the all-exon sequence of one species *S* contains no residues matching an exon *E* found in an all-exon sequence of another species *T*, it suggests that the exon observed in *T* may be present (but unobserved) in *S. Diviner* takes the sequence for the exon observed in T and aligns it to the genomic region in *S* bounded by neighboring exons; if a high-scoring full-length match is found, *Diviner* takes this as evidence for the existence of the exon in *S*.

### Data, Compute Environment, and Timing

We evaluated *Diviner* on the data produced by running *Mirage2* on three protein datasets. The first of these protein datasets is the complete set of human, mouse, and rat isoforms from the *UniProtKB*/*TrEMBL* database (17). We selected these species because they are well studied, and we expect the full contingent of exons in their genomes to be relatively well explored. *TrEMBL* contains a total of 321,793 protein sequences (isoforms) for these three species, representing 26,860 gene families. Approximately 38% of the sequences in *TrEMBL* have been assigned a low annotation scores (*i*.*e*., they have not been confirmed as genuine *in vivo* protein products), so we also tested *Diviner* using the curated set of 73,537 human, mouse, and rat protein sequences in the *UniProtKB*/*SwissProt*. Finally, to explore the impact of broadening scope to include less-sequenced species, we built a third protein dataset by adding the 20,870 *UniProtKB* representative ferret sequences to the set of human, mouse, and rat *SwissProt* sequences. All genomic data used in our testing were downloaded through the UCSC *Genome Browser* (18) (human=hg38, mouse=mm39, rat=rn6, and ferret=musFur1). To compare *Diviner* results with existing annotations, GTF-formatted genome annotation files were composited for each of these four species using the RefSeq and RefGene data from Ensembl (19).

All analysis was performed on a system with 2x AMD EPYC 7642 48-core CPUs and 512GB RAM, but with maximum resource consumption limited to 8 threads and 16GB RAM. *Diviner*’s analysis of the human, mouse, and rat *TrEMBL* sequences (provided as a collection of pre-computed *Mirage2* alignments and protein-to-genome mapping data) completed in 142 minutes. *Diviner* completed analysis of the *SwissProt* human, mouse, and rat dataset in 52 minutes and the dataset built by adding representative ferret *SwissProt* sequences in 99 minutes.

### Analysis of *TrEMBL* Isoforms Uncovers 2120 Human, Mouse, and Rat Exons

By mapping and aligning *TrEMBL*’s 321,793 isoforms from human, mouse, and rat, *Diviner* located a total of 31,460 cases in which (i) an exon found in an isoform belonging to one species *T* is not observed in any *TrEMBL* isoform for another species *S*, and (ii) mapping that exon to the exon-bounded window of *S* produces a high-quality alignment. From perspective of *TrEMBL*, these are newly-revealed exons within species *S*. 12,756 of these revealed exons are in the mouse genome, while 9994 are from rat and 8710 are in human.

The set of isoforms in *TrEMBL* is not the sole source of identified exons for these orgnaisms. We compared the 31,460 new exons to exons captured in RefSeq and RefGene data from Ensembl (19), and found that 29,151 of them overlapped, leaving 2309 novel *Diviner*-discovered exons (1161 in rat, 931 in mouse, 217 in human). We compared these to the set of exons predicted by the tool *SGP-2* (20; available for download from the UCSC Genome Browser), and found 189 overlaps. Thus, a final set of 2120 *Diviner*-discovered exons (1012 in rat, 910 in mouse, 198 in human) appear to be “truly novel” – they are predicted by *Diviner* but are not otherwise observed among competing annotation sources.

Without confirmatory sequencing, it is not possible to be certain that these novel exons are used in some unobserved isoform (*i*.*e*. are expressed *in vivo* under some untested condition). Sequence conservation provides some insight into the validity of the predicted exons. We compute percent identity as the fraction of gap-free alignment columns in which the two sequences share the same amino acid character. As shown in Figure 2, the large majority of “truly novel” recovered exons show sequence alignment percent identity over 50% amino acid identity to their most similar homolog. In alignments with this similarity, structure is expected to be highly conserved (21, 22).

**Fig. 2.**
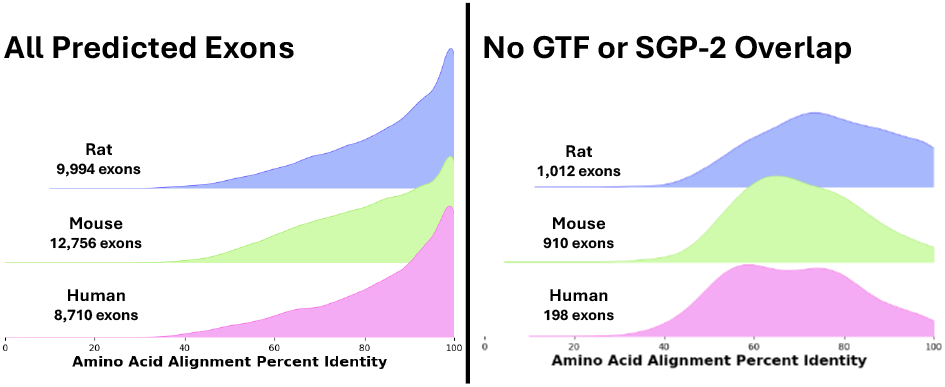
Distributions of alignment percent identity for exons predicted by *Diviner* when analyzing the human, mouse, and rat sequences from the *TrEMBL* dataset, divided according to the species in whose genome the exon was predicted. In cases where multiple query species uncovered the same exon, the pairwise species alignment with the highest percent identity is used.

### Focused Analysis of Curated Proteins in *SwissProt* Reveals 663 High-Confidence Novel Exons

The above analysis based on all *TrEMBL* isoforms is valuable for finding a large set of novel exon candidates, but many of the isoforms in *TrEMBL* are listed as “low quality” and lack experimental support. To establish a more conservative exon set, we also tested *Diviner* on the human, mouse, and rat protein sequences contained in *SwissProt*, which is a curated subset of *TrEMBL* whose constituents can confidently be assumed to represent genuine protein products. *Diviner*’s analysis of the *SwissProt* sequences uncovered a total of 13,177 predicted exon homologs, of which 663 do not overlap with previously annotated or predicted coding regions from RefSeq/RefGene (Figure 3). These 663 high-quality novel exon predictions also show high percent identity, and are found in 516 (2.8%) of the 17,923 gene families in the *SwissProt* dataset.

**Fig. 3.**
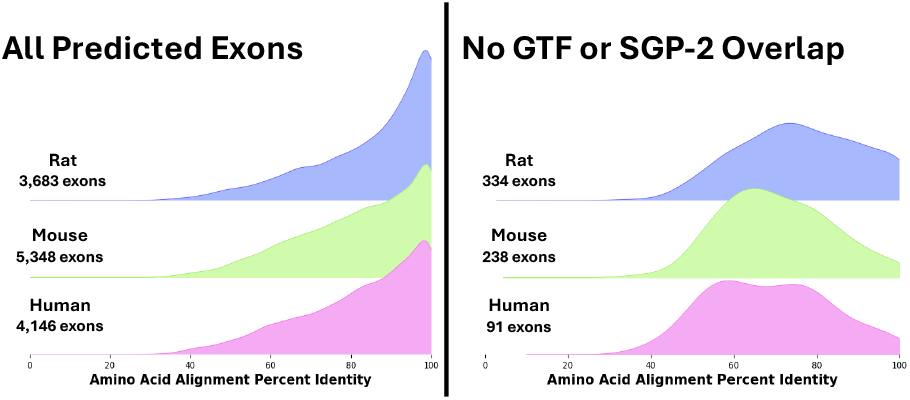
Distributions of alignment percent identity for exons predicted by *Diviner* when analyzing sequences from the *SwissProt* dataset, divided according to the species in whose genome the exon was predicted. In cases where multiple query species uncovered the same exon, the pairwise species alignment with the highest percent identity is used.

### Length and Quality Distributions are Similar Between Predicted and Known Exons

We performed two comparative analyses to assess support for the novel exons predicted by *Diviner*. In the first analysis, we captured exon length and alignment percent identity between *Diviner*-predicted exons that overlapped with GTF-annotated coding regions and “novel” exons that did not overlap with annotated coding regions. As seen in Figure 4, novel exons tend to be shorter in average, while the percent identity of their supporting alignments is within the common range of exons supported by GTF/SGP-2.

**Fig. 4.**
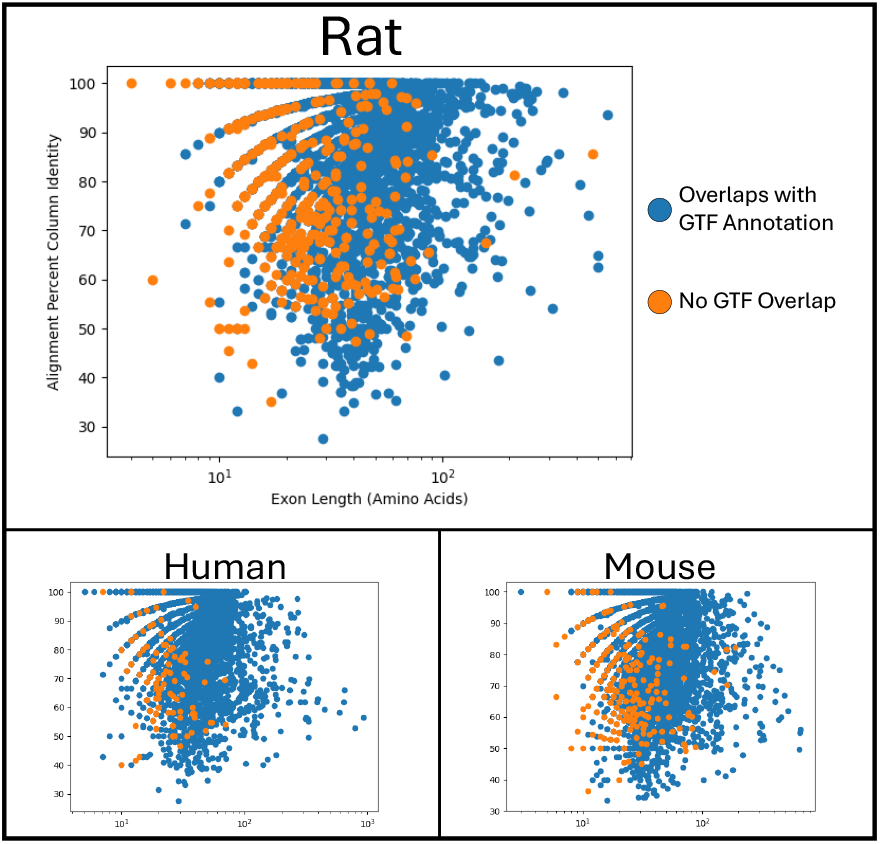
Distributions of alignment percents column identity for exons predicted by *Diviner* when analyzing sequences from the *SwissProt* dataset, divided according to the species in whose genome the exon was predicted. In cases where multiple query species uncovered the same exon, the pairwise species alignment with the highest percent column identity is used.

We also analyzed *Diviner*-predicted exons by comparing their evolutionary divergence (percent identity) to that of other exons within the same gene. For every gene, we computed the alignment percent identity for each exon in that gene’s all-exon pairwise species alignments (Figure 5, see Methods). For each *Diviner*-predicted exon, we computed the difference between (i) the percent identity of its alignment to its most similar query exon and (ii) the average exon alignment percent identity for that species pair and gene family (Figure 6, top row, center column). We also computed the difference in percent identity between the predicted exon and the nearest upstream and downstream exons in the pairwise species alignment (Figure 6, top row, left and right columns, respectively). In all three species, percent identity differences for the full set of *Diviner*-discovered exons are all normal distributions centered near 0%.

**Fig. 5.**
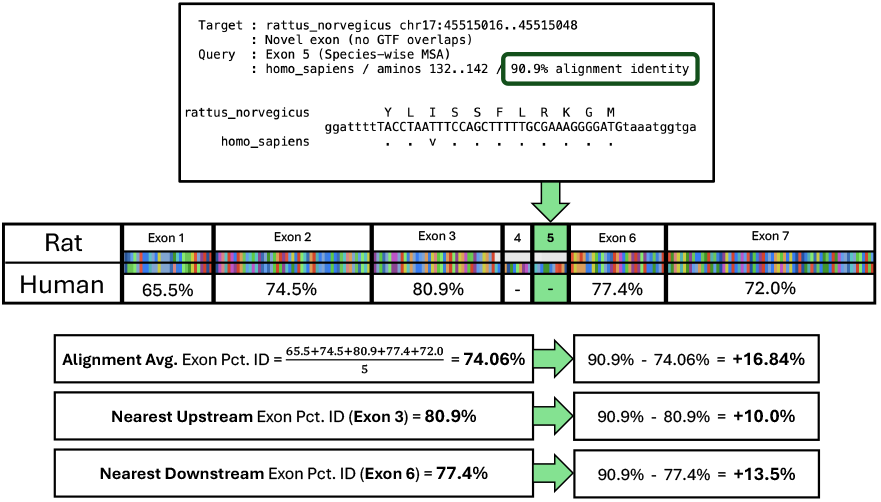
Schematic using *Diviner* ‘s results for the GPX5 gene family to illustrate how reference alignment percentidentity for pairs of species were derived to enable comparison with predicted exons.

**Fig. 6.**
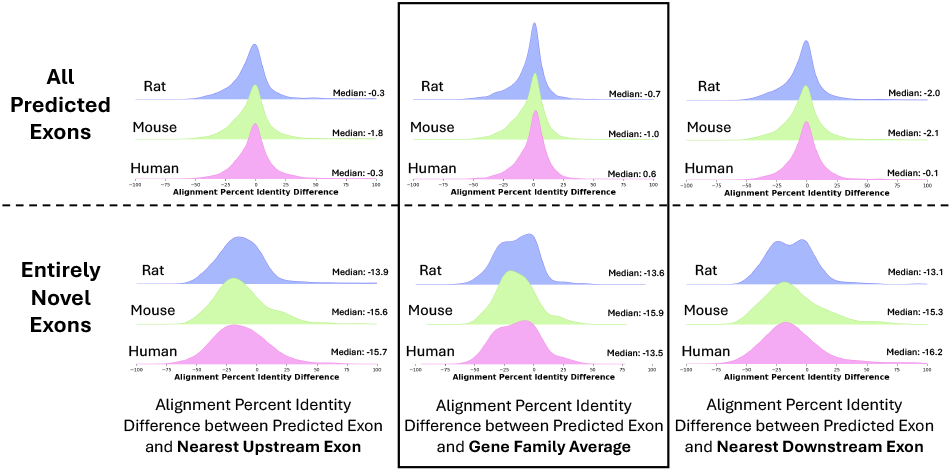
Distributions showing the differences in percents column identity of exons predicted by *Diviner* relative to the percents column identity of exons that are present in the *SwissProt* dataset. The top row represents all exons predicted by *Diviner* (including those that overlap with GTF or *SGP-2* annotations), and the bottom row represents only the “entirely novel” predicted coding regions.

Restricting analysis to entirely-novel exons (not found in GTF or by *SGP-2*), these novel exons show a slight reduction in alignment percent identity relative to other exons for the same gene (Figure 6, bottom row), suggesting that these exons may be under reduced selective pressure relative to more commonly observed exons.

### Distribution of *SwissProt* Annotation Scores for Exons That Suggest Novel Exons

One element of the metadata attached to many *UniProtKB* sequences is an annotation score that numerically indicates the strength of the established evidence that a given amino acid sequence represents a genuine protein product. These scores fall into 5 categories: scores 1 and 2 represent sequences that have been observed at either the protein or transcript level, a score of 3 is assigned to sequences whose production has been inferred by sequence homology, and scores 4 and 5 are assigned to proteins that have only been computationally predicted or are supported by limited evidence. As a class, the coding regions reported by *Diviner* may be thought of as deserving a score of 3 (homology-inferred sequences), but a supplemental piece of information that can be attributed to *Diviner*’s predicted exons is what can be called a “transitive annotation score”: the best annotation score of a known sequence in another species that exhibits an exon-level homology with the predicted exon. The value of computing the transitive annotation scores for predicted exons is that these scores communicate the highest level of evidence supporting a documented homolog of that predicted exon. Strong transitive annotation scores therefore have the potential to elevate our confidence that specific exons predicted by *Diviner* are genuine coding regions.

To assign a transitive annotation score to each *Diviner*predicted exon, we identified the exon sequences in other species that produced an alignment covering at least 85% of the predicted exon’s amino acids. Of these, the exon with the best *UniProtKB* annotation score was identified, and this was recorded as the transitive annotation score of the predicted exon.

Applying this method, a transitive annotation score could be assigned to 74% of *Diviner*-predicted exons from *SwissProt*, and to 80% of the *TrEMBL* sourced exons (Figure 7). The distribution of the transitive annotation scores attributed to exons predicted by *Diviner* is comparable to the distribution of actual annotation scores of *SwissProt* sequences and the large majority of source exons (those used to identify novel exons in other species) have an annotation score of 1. Note that many *SwissProt* sequences have no indicated annotation, while *some* annotation is available for nearly all source exons.

**Fig. 7.**
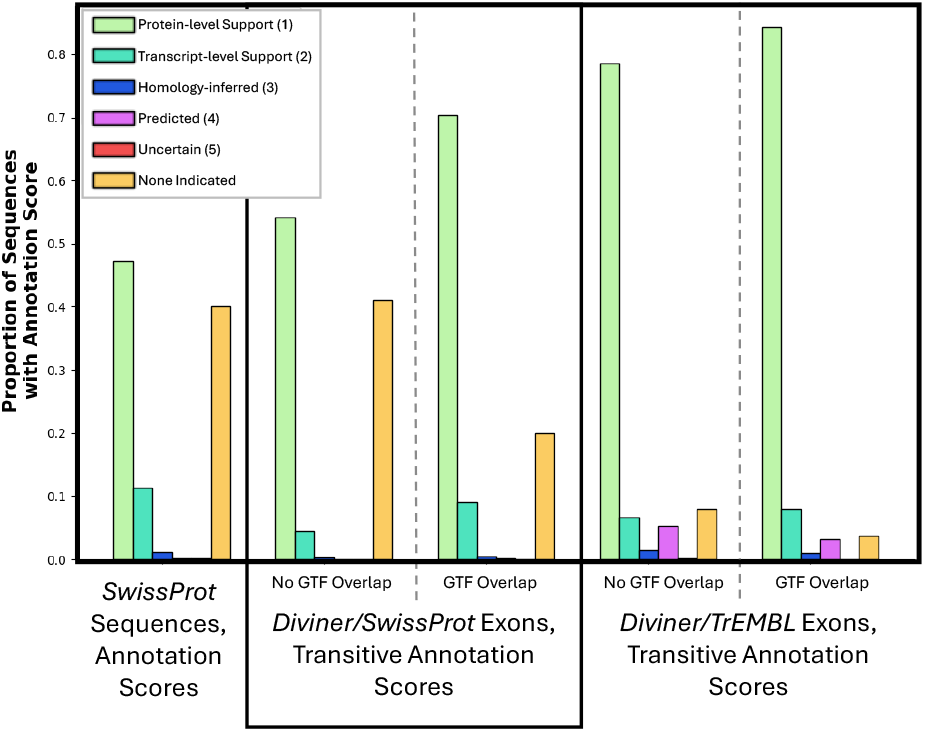
Distributions showing the annotation scores of *SwissProt* sequences compared to the transitive annotation scores attributed to exons predicted by *Diviner*.

### Adding Ferret Sequences Reveals Additional Novel Exons in All Genomes

The above results suggest a strategy for improving the exon annotation in a new genome *G*: add the set of predicted isoforms for *G* to an existing dataset to produce isoform MSAs with *Mirage2*, and use *Diviner* to identify new exons in *G* based on their similarity to observed/known exons in other organisms. As a demonstration of this approach, we ran *Diviner* on a dataset constructed by adding ferret protein sequences from *UniProtKB* (musFur1) to the collection of human, mouse, and rat sequences from *SwissProt*.

### New exons in ferret based on other organisms

As anticipated, *Diviner* uncovered many ferret exons (6470) that are not found in the ferret proteome available from *UniProtKB*. Of these, 699 do not overlap with GTF-annotated coding regions. The UCSC *Genome Browser* contains no *SGP-2* predictions for ferret. Most of these newly-identified exons are supported by exons in only one (610) or two (68) of the human/mouse/rat isoform sets, highlighting the notion that different conditions have been explored by sequencing efforts on the various organisms.

### New exons in other organisms based on ferret

Importantly, the *Diviner* analysis also discovered new exons in the other species that were not present in their *SwissProt* isoforms: 6872 human exons, 7017 mouse exons, and 4398 rat exons (Figure 8, left). Of these, 570 are entirely novel exon predictions (not previously present in GTF or *SGP-2* annotations, nor in the earlier mouse-rat-human *Diviner* run) that were only located with the introduction of ferret sequences: 335 in human, 136 in mouse, and 99 in rat.

**Fig. 8.**
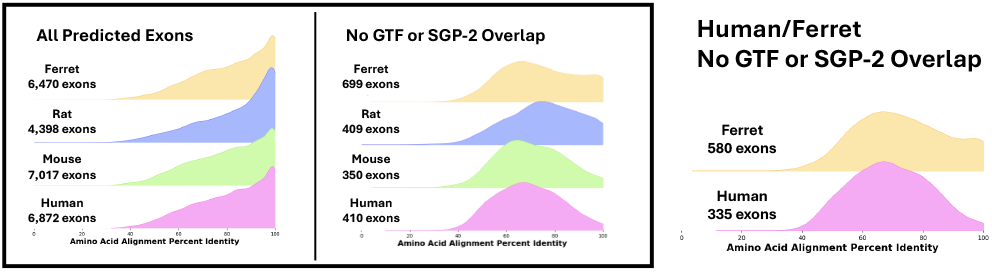
Left Box: The distribution of alignment percents column identity for exons predicted by *Diviner* when analyzing the dataset constructed by adding representative ferret sequences to the *SwissProt* human, mouse, and rat sequences. Center and Right: Distributions of alignment percent column identity for exons predicted by *Diviner* when reduced to results that would be achieved by searching only with ferret and human sequences as inputs.

While these predicted exons are new to mouse+rat+human sequence in *SwissProt* and do not overlap with GTF annotated or *SGP-2* predicted coding regions, they could possibly overlap with exons that are present in the *TrEMBL* dataset. To determine whether the ferret-informed exons found in *SwissProt* analysis were also discoverable using *TrEMBL* sequences, we compared the two sets (*SwissProt*/ferret+mouse+rat+human exons vs *TrEMBL*/mouse+rat+human exons). Only 8 of the “novel” exons (7 human and 1 mouse) found during the *SwissProt*/ferret+mouse+rat+human search are already present in *TrEMBL*. Another 103 *SwissProt*/ferret+mouse+rat+human exons could alternatively have been identified with *TrEMBL*/mouse+rat+human analysis (17 human, 44 mouse, 42 rat). The remaining 459 (311 human, 91 mouse, 57 rat) were found to be unique to *SwissProt* searches with ferret sequences (see Figure 9).

**Fig. 9.**
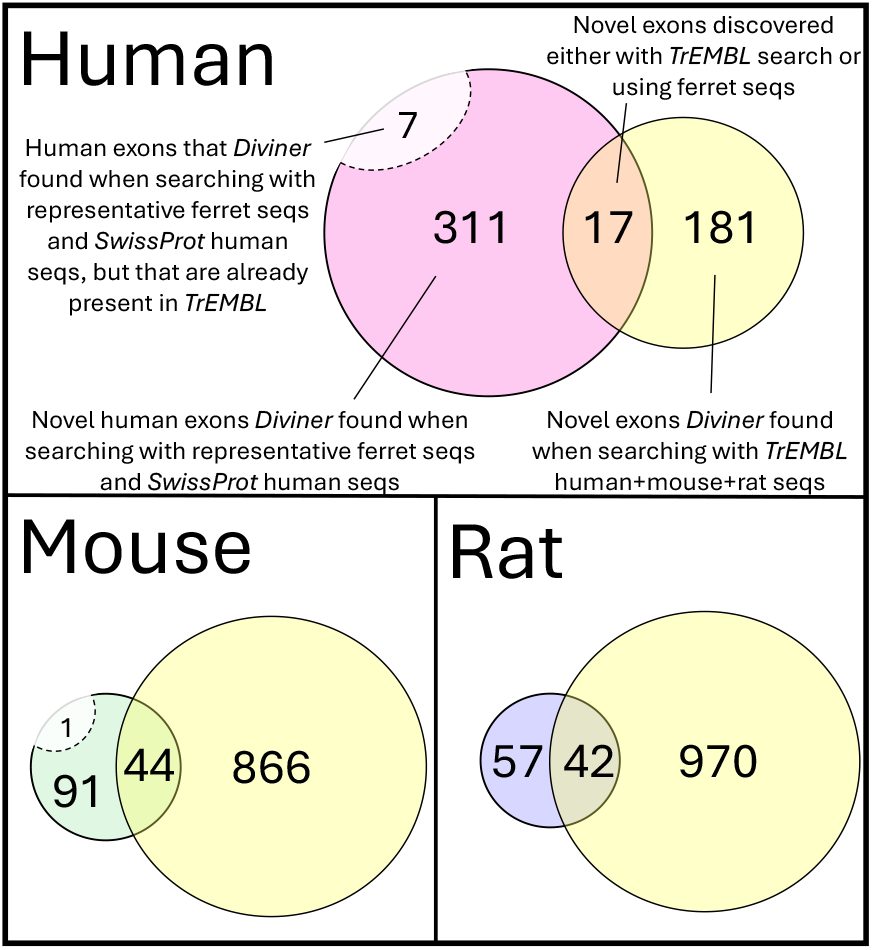
Results of cross-tabulating novel exons (exons without GTF or *SGP-2* overlap) found during *Diviner* ‘s analysis of the dataset combining human, mouse, and rat *SwissProt* sequences with the representative proteome for ferret (removing exons that overlapped with exons present in *TrEMBL*) and the human, mouse, and rat *TrEMBL* dataset.

### A Predicted Rat Exon Produces a Similar Structure to a Known Human Isoform

Consider a newly-discovered exon in a protein-coding gene in species *X*, supported by an isoform from species *Y* . The exon in *X* will, if expressed, be incorporated into a larger encoded protein; there, it is expected to produce a three-dimensional structure that is similar to the structure of the source protein from *Y* . To explore this expectation, we identified a “novel” exon in the rat UGT2A1 gene that is supported by a human exon in the homolgous gene. The predicted exon is 210 amino acids long (a large fraction of the 527 amino acid length of the known rat isoform), shares an 81.4% alignment identity between rat and human exons, and does not overlap with any annotated or predicted coding regions on the rat genome (Figure 10, left). Additionally, relative to the primary rat UGT2A1 isoform in *SwissProt*, the human isoform simply adds the matching exon into the sequence without making any other modifications to the primary isoform. This allows a simple test: computationally “splice” the novel rat UGT2A1 exon into the primary rat isoform from *SwissProt*, and evaluate the impact on predicted structure.

**Fig. 10.**
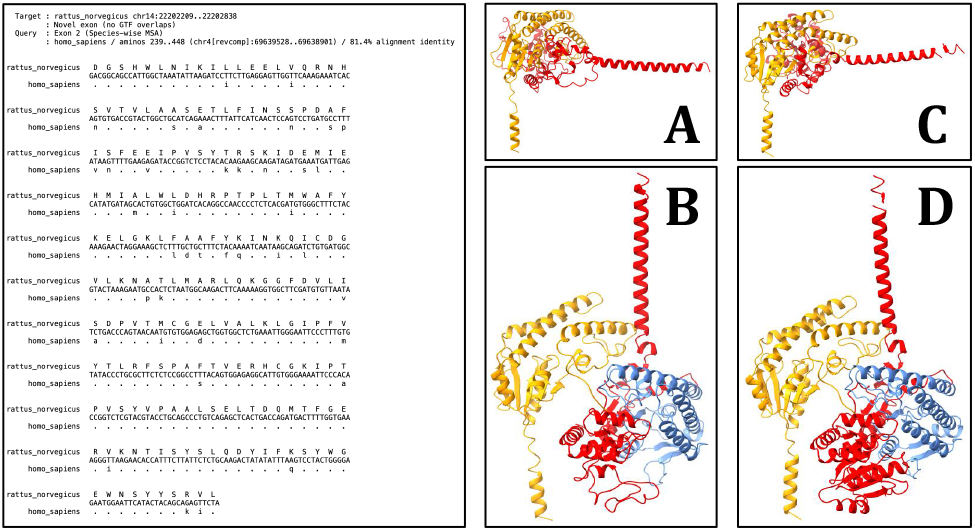
Left: *Diviner* ‘s prediction of a novel UGT2A1 exon in the rat genome, located by virtue of its homology to a known human UGT2A1 exon. Right: *ChimeraX* visualizations of the predicted structures for: [A] the primary human UGT2A1 isoform in *SwissProt* (accession: P0DTE4), [B] an alternative human UGT2A1 isoform in *SwissProt* (accession: P0DTE4-4), containing the exon that served as the query for the pictured *Diviner* search result, [C] the sole rat UGT2A1 isoform in *SwissProt* (accession: P36510), and [D] the protein sequence produced by incorporating the novel rat exon predicted by *Diviner* into the rat UGT2A1 isoform pictured in [C]. The rat exon predicted by *Diviner* and its human homolog are colored blue. The N-terminal portion of each protein is colored yellow and the C-terminal portion is colored red.

We considered predicted structure of four protein sequences: the primary human and rat isoforms, the human isoform whose exon served as the query for the search that uncovered the novel rat exon, and the simulated rat isoform that we produced by adding the predicted exon to the primary rat isoform. For protein structure prediction, we used *ESMFold* (23) on each sequence (default parameters), selected specifically because it predicts structure from an individual sequence.

Predicted structures were visualized using *ChimeraX* (24). The primary UGT2A1 isoforms (absent the novel exon) for human (Figure 10A) and rat (Figure 10C) exhibit nearly identical three-dimensional structures. The alternative human isoform (Figure 10B) produces a structure that is distinct from the primary isoform due to its inclusion of the additional exon (colored blue in the figure); the simulated rat isoform (Figure 10D) produces a three-dimensional structure that is highly similar to the structure of the matching human isoform, lending support to the belief that the alternative rat isoform is also produced as a genuine functional protein.

## Data Availability

The collection of *Diviner*-discovered exons is available for download at https://zenodo.org/doi/10.5281/zenodo.11116253. New exon sets are provided for the human, mouse, rat, and ferret genomes. Each set contains the list of new exons, an indicator of novelty relative to GTF/SGP-2, and information about the supporting exon and alignment.

## Methods

*Diviner* identifies previously-undocumented exons by following four steps: (1) produce an inter-species multiple sequence alignment (MSA) of all provided isoforms, respecting exons boundaries; (2) for each species, reduce all isoforms to a single *all-exon* representative sequence, yielding a compact MSA with one sequence per species; (3) using this all-exons MSA, identify alignment gaps that suggest exons missing in the sequence for one species but present in another; then (4) search for the missing exon in the genomic range bounded by neighboring exons.

### Producing Species-Wise All-Exon Alignments

Let *S* = (*s*_1_, *s*_2_, …, *s*_*n*_) be the set of *n* species represented in the input, and let *I*_*g*_ be the complete set of available protein isoforms for a specific gene *g*, with *I*_*g*_(*k*) being the subset of those isoforms associated with species *s*_*k*_. Before running *Diviner*, a gene-specific MSA of all sequences in *I*_*g*_ must be produced using our *Mirage2* software package (16). *Mirage2* produces gene-specific isoform MSAs by first mapping each sequence in a set *I*_*g*_(*k*) to the appropriate reference genome, then using those nucleotide mapping coordinates as the basis for intra-species sequence alignment in which all isoforms in *I*_*g*_(*k*) are organized such that amino acids mapping to the same genomic position are placed in the same column. This procedure ensures that each intra-species alignment captures and adheres to splice boundaries. Inter-species alignment is performed by merging (aligning) these intra-species MSAs following a profile-based Needleman-Wunsch (25) implementation with a modifier to encourage maintenance of splice boundaries across species (for all Mirage details, see 16, 26). *Mirage2* provides two products that are used by *Diviner*. The primary product is an amino acid multiple sequence alignment *M*_*g*_ of all isoforms of gene *g*. In *M*_*g*_, each row *i* corresponds to one isoform sequence from *I*_*g*_, and the sequences are spread out so that related amino acids share a column *j*: *M*_*g*_(*i, j*) contains a letter if an amino acid from sequence *i* is associated with alignment column *j*, and otherwise contains a ‘-’. The secondary product from *Mirage2* is a mapping matrix *P*_*g*_ that stores, for each aligned amino acid in *M*_*g*_, the position of the middle nucleotide of the encoding codon; if *M*_*g*_(*i, j*) is a gap-character, *P*_*g*_(*i, j*) == *NU LL* .

Rather than consider all isoform sequences for a gene *g* independently, *Diviner* collapses isoforms to produce a single sequence *E*_*g*_(*k*) per species *s*_*k*_, corresponding to the union of all exons observed for gene *g* for *s*_*k*_, ordered by their position in the genome. For each species, we call the collapsed set the *all-exon representation*. By construction, *E*_*g*_(*k*) will not necessarily correspond to any individual isoform. The resulting collapsed alignment matrix *C* contains a single row *C*_*g*_(*k*) for each species *k*. The collapsed matrix *C*_*g*_ is the same length as *M*_*g*_, and contains gap characters (‘-’) where no sequence in *I*_*g*_(*k*) contained an amino acid. *C*_*g*_ and is complemented by a codon position matrix *Q*_*g*_ that plays a role analagous to *P*_*g*_, but for the all-exon alignment. Usually, the task of collapsing all isoforms of a gene/species pair is straightforward, because a codon used in one isoform will be either used (encoding the same amino acid) or not used in other isoforms. Occasionally, a coding region may code in more than one frame in different alternative splicing products (26); in this case, a simple majority-rule vote is used to determine the consensus amino acid and nucleotide coordinate for each species at each column in *E*_*g*_ and *Q*_*g*_.

Finally, for an alignment *C*_*g*_ containing *f* exon blocks, *Diviner* identifies a set *X* of tuples {(1, *x*_1_), (*x*_1_ + 1, *x*_2_), (*x*_2_ + 1, *x*_3_), …, (*x*_*f*−1_ + 1, *m*)} that divides the alignment into adjacent column ranges corresponding to the exon boundaries. These enable *Diviner* to easily detect where an exon present in one species is lacking a homolog in another species.

### Identifying Candidate “Missing” Exons

To identify missing-exon candidates, *Diviner* scans through each row in *E*_*g*_ and checks, for each range tuple in *X*, if the corresponding string in *C*_*g*_ consists entirely of gap characters. Suppose such a range is identified in row *k*, beginning at column *b* and ending at column *e*: this indicates that at least one of the other species *s*_*i*_, *i k* has an isoform that contains an exon represented in range (*b*..*e*), but species *s*_*k*_ does not (Figure 11).

**Fig. 11.**
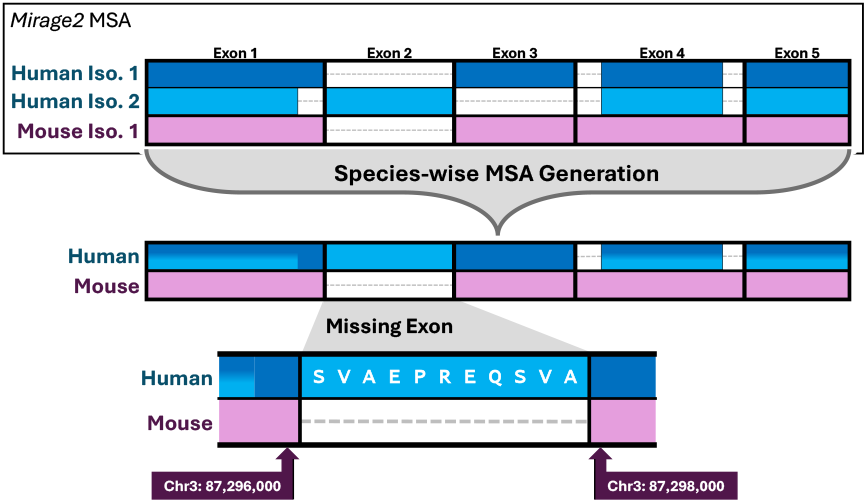
*Diviner* identifies windows of genomic sequence where a “missing” exon would be expected to reside by deriving a species-wise spliced alignment (and corresponding protein-to-genome mapping coordinates) from a *Mirage2* multiple sequence alignment and then extracting relevant sequence data around exons that have no homolog in one or more other species.

### Locating Missing Exons

When an exon is found to be missing from the isoforms of species *s*_*k*_, it is either actually not present in the genome of *s*_*k*_, or has simply been unobserved among the sampled isoforms. Diviner performs a localized sequence alignment to identify a potential location of the unobserved exon (Figure 12).

**Fig. 12.**
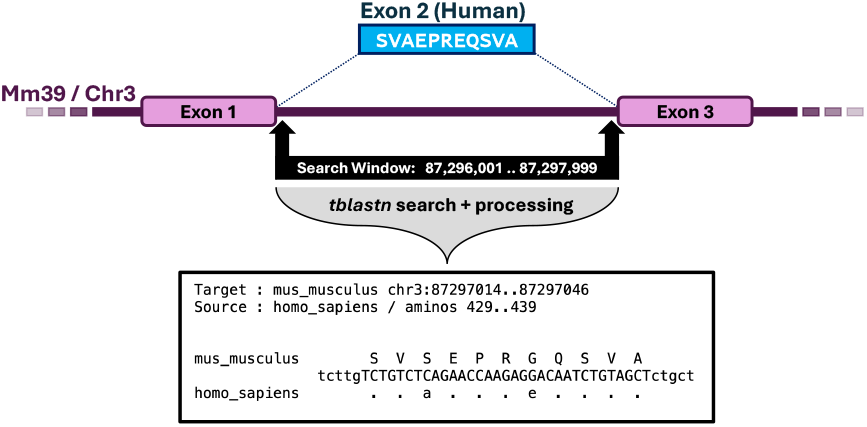
Once *Diviner* has identified a region of the species-wise alignment that suggests an exon is missing a homolog in another species, it extracts the region of that species’ genome where that exon homolog is expected to reside and uses *tblastn* to search for the missing homolog. If a strong match is found, *Diviner* extracts the genomic sequence corresponding to the match and produces an alignment of the known exon to its predicted homolog.

Since (*b*..*e*) is the column range for the exon missing from species *s*_*k*_, position *b*− 1 contains the last amino acid upstream of the missing exon. Let *L* = *Q*_*g*_(*k, b* − 1) capture the center nucleotide of that amino acid. This is the left-most position that the missing exon could be found. Similarly, let *R* = *Q*_*g*_(*k, e* + 1) capture the start of the nearest downstream exon. In the edge case where there is no upstream exon, *Diviner* sets *L* = *R* − 25, 000; conversely, if there is no downstream exon, *R* = *L* + 25, 000. The resulting range, *L*..*R*, defines the range in which the missing exon can be expected to reside. (In all cases here, notation assumes the protein is encoded on the forward-strand; reverse strand positions are trivially adjusted to account for inverted order).

To construct its amino acid query sequence(s), *Diviner* captures the amino acids for each candidate *other* sequence *O* = *C*_*g*_(*i, b*)..*C*_*g*_(*i, e*), *i* ≠ *k*. If *O* contains at least 6 amino acids, *Diviner* uses the fast translated search tool *tblastn* (27) to search *O* against the search window *L*..*R* on the genome of species *s*_*k*_, and records the highest-scoring non-overlapping hits. Each of these *tblastn* hits is treated as the seed of a putative exon and serves as the basis of re-alignment using an implementation of the Smith-Waterman local alignment algorithm (28). The resulting alignments of each query exon to the target region are recorded for output.

### Formatted Output for Predicted Exons

*Diviner* reports data about its predicted exons in several formats. The most straightforward representations of the exons predicted by *Diviner* are recorded as a set of .bed files (one per target species) (Figure 13). Each line in these .bed files communicates a specific nucleotide range on the species’ reference genome corresponding to a predicted exon. Information about the gene family and exon indices of the query sequence are also provided, enabling easy attribution and manual examination of the region of the species-wise MSA where the newly predicted exon would reside. The alignment percent column identity for the predicted exon is also included as the penultimate field of each line.

**Fig. 13.**
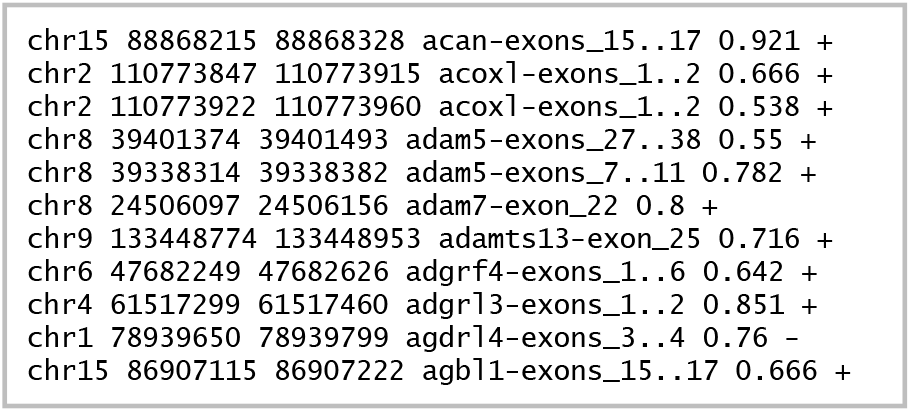
Example of a .bed file produced by *Diviner*, listing exons uncovered in the human genome.

*Diviner* also produces an output format designed for improved exploration of exon candidates: a file named ‘Hits-byPct-ID.out’ sorts the entire collection of predicted exons in descending order of alignment percent column identity (Figure 14). Additionally, this file communicates the region on the target genome corresponding to the predicted exon, the amino acid length of the exon, and whether the predicted exon overlaps with a GTF-annotated coding region.

**Fig. 14.**
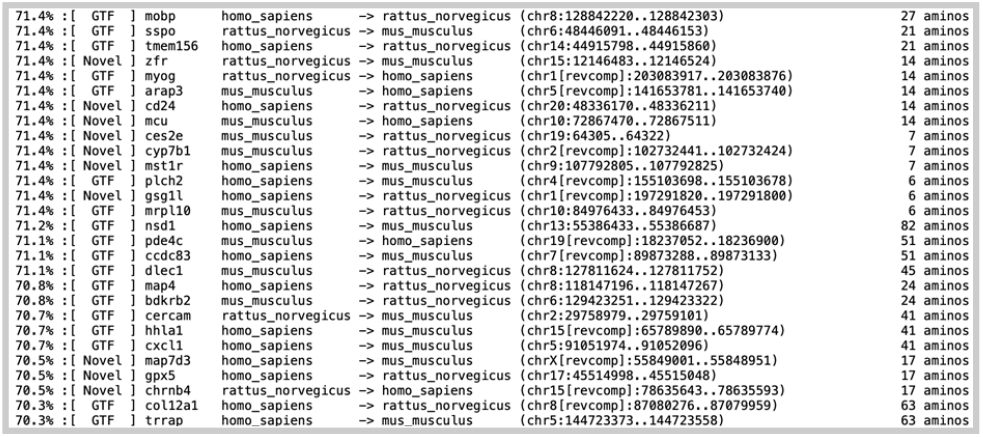
A portion of the Hits-by-Pct-ID.out file produced by *Diviner* when run on the *SwissProt* dataset of human, mouse, and rat isoforms.

Finally, *Diviner*’s most detailed result data are broken down by gene family in a folder called ‘Results-by-Gene’. The subdirectories of this folder contain descriptive alignment files that provide visual representations of the alignment of each predicted exon to its query sequences along with the coding locations of those query exons on their species’ genomes (Figure 15). Further, *Diviner* preserves the *C* and *Q* data for each family (*C* in an aligned-FASTA-formatted file and *Q* in a file that uses the same format as *Mirage2*’s mapping output files), simplifying the work needed to write programs for output verification and downstream analyses.

**Fig. 15.**
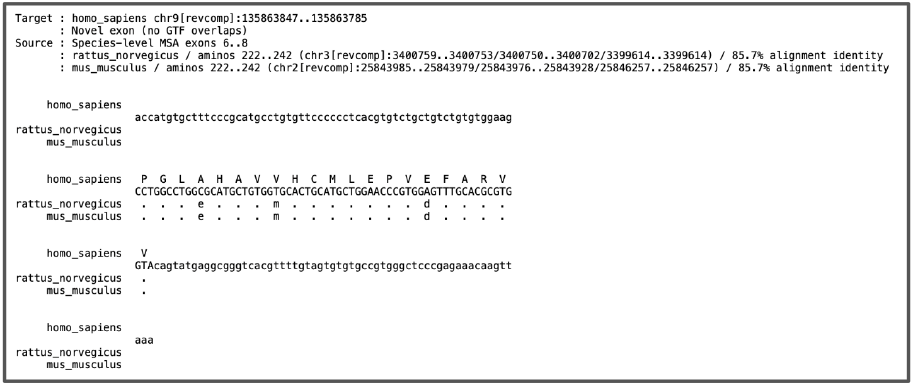
An example of a file produced by *Diviner* displaying an alignment of known mouse and rat CAMSAP1 exons to a novel exon predicted in the human genome.

## Discussion

### Existence of Exon Homologs May Bolster Confidence for Predicted or Uncertain Isoforms

32,978 of the human, mouse, and rat isoforms contained in the *TrEMBL* dataset have an annotation score that indicates they are either computationally predicted or are supported by uncertain evidence. Among these low-scoring isoforms, there are sequences that can be characterized by their use of one or more exons that are not included in any of the better-supported sequences in their gene family a sequence’s low score may be attributed to its use of a unique and unverified exon. One method that could be employed to obtain further support for these low-scoring isoforms involves using *Diviner* to search for coding regions on other species’ genomes that exhibit strong homology to these unverified characteristic exons. If there is evidence that the characteristic exons of low-scoring isoforms have been evolutionarily conserved in other organisms, that could be interpreted as a source of additional support for their genuine *in vivo* production.

### Sketching a Pipeline for Using *Diviner* in *ab initio* Exon Prediction

The results that we achieved by using *Diviner* to uncover novel exons in the ferret genome indicate that it could be a useful tool for predicting coding regions in other species that have not been extensively studied. As sequencing entire genomes becomes more affordable, *Diviner* could be used to locate exons on entirely unannotated genomes. The challenge for using *Diviner* on entirely unannotated genomes is that *Diviner* requires access to at least one protein per gene that can provide the genomic context in which additional exons can be identified. For species that do not have a reference proteome (or may not have reference sequences available in all gene families), it is therefore necessary to derive at least one amino acid sequence that can be mapped to the species’ genome to anchor *Diviner*’s search for exon homologs.

To acquire an amino acid sequence that can direct *Diviner* towards the appropriate search region for a particular gene family on a novel genome, we propose the following sketch of a pipeline: (i) for each gene family of interest, acquire a protein sequence belonging to that family from another species (*e*.*g*., its primary human isoform in *SwissProt*); (ii) produce a spliced alignment using a tool such as *miniprot* (15); (iii) treat the translated sequences of any regions on the target genome that align well to the query protein as a set of “seed” exons, and concatenate these exons into a “reference” protein sequence for that gene family in the novel species. Following alignment of this synthetic protein sequence to the actual proteins of other species by *Mirage2, Diviner* would have the input data required to search for additional exon homologs in the correct region of the novel genome.

## ACKNOWLEDGEMENTS

We gratefully acknowledge the computational resources and expert administration provided by the University of Montana’s Griz Shared Computing Cluster (GSCC) and high performance computing (HPC) resources supported by the University of Arizona TRIF, UITS, and Research, Innovation, and Impact (RII) and maintained by the UArizona Research Technologies department. AJN and TJW were supported by NIH NIGMS R01GM132600 and NHGRI R21HG012283.

## Bibliography

1. Panagiotis Papasaikas and Juan Valcárcel. The spliceosome: the ultimate RNA chaperone and sculptor. Trends in biochemical sciences, 41(1):33–45, 2016.

2. Robert J Weatheritt, Norman E Davey, and Toby J Gibson. Linear motifs confer functional diversity onto splice variants. Nucleic acids research, 40(15):7123–7131, 2012.

3. Olga Kelemen, Paolo Convertini, Zhaiyi Zhang, Yuan Wen, Manli Shen, Marina Falaleeva, and Stefan Stamm. Function of alternative splicing. Gene, 514(1):1–30, 2013.

4. Eric T Wang, Rickard Sandberg, Shujun Luo, Irina Khrebtukova, Lu Zhang, Christine Mayr, Stephen F Kingsmore, Gary P Schroth, and Christopher B Burge. Alternative isoform regulation in human tissue transcriptomes. Nature, 456(7221):470–476, 2008.

5. Markus J Sommer, Sooyoung Cha, Ales Varabyou, Natalia Rincon, Sukhwan Park, Ilia Minkin, Mihaela Pertea, Martin Steinegger, and Steven L Salzberg. Structure-guided isoform identification for the human transcriptome. Elife, 11:e82556, 2022.

6. Kuo-Feng Tung, Chao-Yu Pan, Chao-Hsin Chen, and Wen-chang Lin. Top-ranked expressed gene transcripts of human protein-coding genes investigated with GTEx dataset. Scientific Reports, 10(1):16245, 2020.

7. Nicolas Scalzitti, Anne Jeannin-Girardon, Pierre Collet, Olivier Poch, and Julie D Thompson. A benchmark study of ab initio gene prediction methods in diverse eukaryotic organisms. BMC genomics, 21:1–20, 2020.

8. Annotation score - UniProt. https://www.uniprot.org/help/annotation_score. xAccessed: Apr 5, 2024.

9. Nicholas T Ingolia, Gloria A Brar, Silvia Rouskin, Anna M McGeachy, and Jonathan S Weissman. The ribosome profiling strategy for monitoring translation in vivo by deep sequencing of ribosome-protected mrna fragments. Nature protocols, 7(8):1534–1550, 2012.

10. Gloria A Brar and Jonathan S Weissman. Ribosome profiling reveals the what, when, where and how of protein synthesis. Nature reviews Molecular cell biology, 16(11):651–664, 2015.

11. Jens Keilwagen, Michael Wenk, Jessica L Erickson, Martin H Schattat, Jan Grau, and Frank Hartung. Using intron position conservation for homology-based gene prediction. Nucleic acids research, 44(9):e89–e89, 2016.

12. Ning Yu, Zeng Yu, Bing Li, Feng Gu, and Yi Pan. A comprehensive review of emerging computational methods for gene identification. Journal of Information Processing Systems, 12(1), 2016.

13. Alexandre Lomsadze, Paul D Burns, and Mark Borodovsky. Integration of mapped RNA-Seq reads into automatic training of eukaryotic gene finding algorithm. Nucleic acids research, 42(15):e119–e119, 2014.

14. Hiroaki Iwata and Osamu Gotoh. Benchmarking spliced alignment programs including Spaln2, an extended version of Spaln that incorporates additional species-specific features. Nucleic acids research, 40(20):e161–e161, 2012.

15. Heng Li. Protein-to-genome alignment with miniprot. Bioinformatics, 39(1):btad014, 2023.

16. Alexander J Nord and Travis J Wheeler. Mirage2’s high-quality spliced protein-to-genome mappings produce accurate multiple-sequence alignments of isoforms. Plos one, 18(5): e0285225, 2023.

17. Alex Bateman, Maria-Jesus Martin, Sandra Orchard, Michele Magrane, Shadab Ahmad, Emanuele Alpi, Emily H Bowler-Barnett, Ramona Britto, Hema Bye-A-Jee, Austra Cukura, et al. UniProt: the universal protein knowledgebase in 2023. Nucleic Acids Research, 51 (D1), 2022.

18. Luis R Nassar, Galt P Barber, Anna Benet-Pagès, Jonathan Casper, Hiram Clawson, Mark Diekhans, Clay Fischer, Jairo Navarro Gonzalez, Angie S Hinrichs, Brian T Lee, et al. The UCSC genome browser database: 2023 update. Nucleic acids research, 51(D1):D1188–D1195, 2023.

19. Fergal J Martin, M Ridwan Amode, Alisha Aneja, Olanrewaju Austine-Orimoloye, Andrey G Azov, If Barnes, Arne Becker, Ruth Bennett, Andrew Berry, Jyothish Bhai, et al. Ensembl 2023. Nucleic acids research, 51(D1):D933–D941, 2023.

20. Roderic Guigo, Emmanouil T Dermitzakis, Pankaj Agarwal, Chris P Ponting, Genís Parra, Alexandre Reymond, Josep F Abril, Evan Keibler, Robert Lyle, Catherine Ucla, et al. Comparison of mouse and human genomes followed by experimental verification yields an estimated 1,019 additional genes. Proceedings of the National Academy of Sciences, 100(3): 1140–1145, 2003.

21. Gajendra PS Raghava and Geoffrey J Barton. Quantification of the variation in percentage identity for protein sequence alignments. BMC bioinformatics, 7:1–4, 2006.

22. Patrice Koehl and Michael Levitt. Sequence variations within protein families are linearly related to structural variations. Journal of molecular biology, 323(3):551–562, 2002.

23. Jeliazko R Jeliazkov, Diego del Alamo, and Joel D Karpiak. ESMfold hallucinates native-like protein sequences. bioRxiv, pages 2023–05, 2023.

24. Elaine C Meng, Thomas D Goddard, Eric F Pettersen, Greg S Couch, Zach J Pearson, John H Morris, and Thomas E Ferrin. UCSF ChimeraX: Tools for structure building and analysis. Protein Science, 32(11):e4792, 2023.

25. Saul B Needleman and Christian D Wunsch. A general method applicable to the search for similarities in the amino acid sequence of two proteins. Journal of molecular biology, 48(3): 443–453, 1970.

26. Alex Nord, Peter Hornbeck, Kaitlin Carey, and Travis Wheeler. Splice-aware multiple sequence alignment of protein isoforms. In Proceedings of the 2018 ACM International Conference on Bioinformatics, Computational Biology, and Health Informatics, pages 200–210, 2018.

27. E Michael Gertz, Yi-Kuo Yu, Richa Agarwala, Alejandro A Schäffer, and Stephen F Altschul. Composition-based statistics and translated nucleotide searches: improving the TBLASTN module of BLAST. BMC biology, 4(1):1–14, 2006.

28. Temple F Smith, Michael S Waterman, et al. Identification of common molecular subsequences. Journal of molecular biology, 147(1):195–197, 1981.

